# Contrastive Learning Enables Epitope Overlap Predictions for Targeted Antibody Discovery

**DOI:** 10.1101/2025.02.25.640114

**Authors:** Clinton M. Holt, Alexis K. Janke, Parastoo Amlashi, Parker J. Jamieson, Toma M. Marinov, Ivelin S. Georgiev

## Abstract

Computational epitope prediction remains an unmet need for therapeutic antibody development. We present three complementary approaches for predicting epitope relationships from antibody amino acid sequences. First, we analyze ∼18 million antibody pairs targeting ∼250 protein families and establish that a threshold of >70% CDRH3 sequence identity among antibodies sharing both heavy and light chain V-genes reliably predicts overlapping-epitope antibody pairs. Next, we develop a supervised contrastive fine-tuning framework for antibody large language models which results in embeddings that better correlate with epitope information than those from pre-trained models. Applying this contrastive learning approach to SARS-CoV-2 receptor binding domain antibodies, we achieve 82.7% balanced accuracy in distinguishing same-epitope versus different-epitope antibody pairs and demonstrate the ability to predict relative levels of structural overlap from learning on functional epitope bins (Spearman *ρ* = 0.25). Finally, we create AbLang-PDB, a generalized model for predicting overlapping-epitope antibodies for a broad range of protein families. AbLang-PDB achieves five-fold improvement in average precision for predicting overlapping-epitope antibody pairs compared to sequence-based methods, and effectively predicts the amount of epitope overlap among overlapping-epitope pairs (*ρ* = 0.81). In an antibody discovery campaign searching for overlapping-epitope antibodies to the HIV-1 broadly neutralizing antibody 8ANC195, 70% of computationally selected candidates demonstrated HIV-1 specificity, with 50% showing competitive binding with 8ANC195.

Together, the computational models presented here provide powerful tools for epitope-targeted antibody discovery, while demonstrating the efficacy of contrastive learning for improving epitope-representation.

## Introduction

Monoclonal antibody therapeutics have revolutionized modern medicine since their first FDA approval in 1986, with blockbuster treatments for cancers, autoimmune diseases, and infectious diseases generating billions in annual revenue^1^. Beyond therapeutics, antibodies serve as fundamental research tools and provide crucial insights into immune responses to vaccines and pathogens. Despite their clinical success, developing therapeutic antibodies remains resource-intensive, with epitope characterization—identifying the specific region on an antigen where an antibody binds—posing a significant bottleneck^2^. For example, in the development of broadly neutralizing antibodies against HIV-1, epitope mapping is critical to ensuring efficacy across diverse viral strains^3^.

Epitope characterization typically proceeds through three complementary approaches:

(1) structural mapping to define physical contact points between antibody and antigen, (2) functional mapping to identify binding-critical residues through mutation, and (3) competition binding experiments to group antibodies that interfere with each other’s binding. Each approach helps guide therapeutic development, whether identifying sites of vulnerability on pathogens or developing complementary antibody combinations ^4–6^.

Understanding the similarities and differences (or the level of overlap) between the epitopes of different antibody candidates provides critical information that can be utilized when developing antibody therapeutics. For example, in pandemic response efforts against a newly emerging virus, the selection of two or more non-competing antibodies which synergize to form a more effective drug than either individual antibody can be critical for counteracting potential virus escape. In other cases, identifying multiple antibodies against the same functionally important epitope can provide a larger set of candidates for further evaluation, down-selection, and development.

While experimental approaches for antibody epitope characterization are undoubtedly effective, computational approaches can present an efficient and cost-effective alternative. Generally, computational approaches can interrogate the relationship between antibody sequence features and epitope similarity, in order to predict the level of epitope overlap between antibody candidates (Fig. 1). These approaches range from direct comparisons of the full amino acid sequence or just the complementarity determining region 3 (CDR3) amino acid sequence within gene groups, to comparing predicted structures or predicted antigen-binding residues^5,7–16^. While the direct sequence-based methods have shown success in clustering functionally-related antibodies, the antibody sequence similarity thresholds utilized by these approaches have been rigorously validated for only a few antigens and epitopes ^5,8–10,17^. The indirect approaches allow for searching a broader antibody sequence-space, but accuracies are low and are unable to detect overlapping epitope antibodies using distinct structural mechanisms, such as targeting the same site from different angles—an aspect that can significantly influence Fc effector functions and binding breadth ^16,18–20^. This limitation is particularly problematic when searching for therapeutic candidates, where expanding the candidate pool beyond highly similar structures could be necessary to overcome challenges like low yields or suboptimal binding properties ^21^.

**Figure 1:**
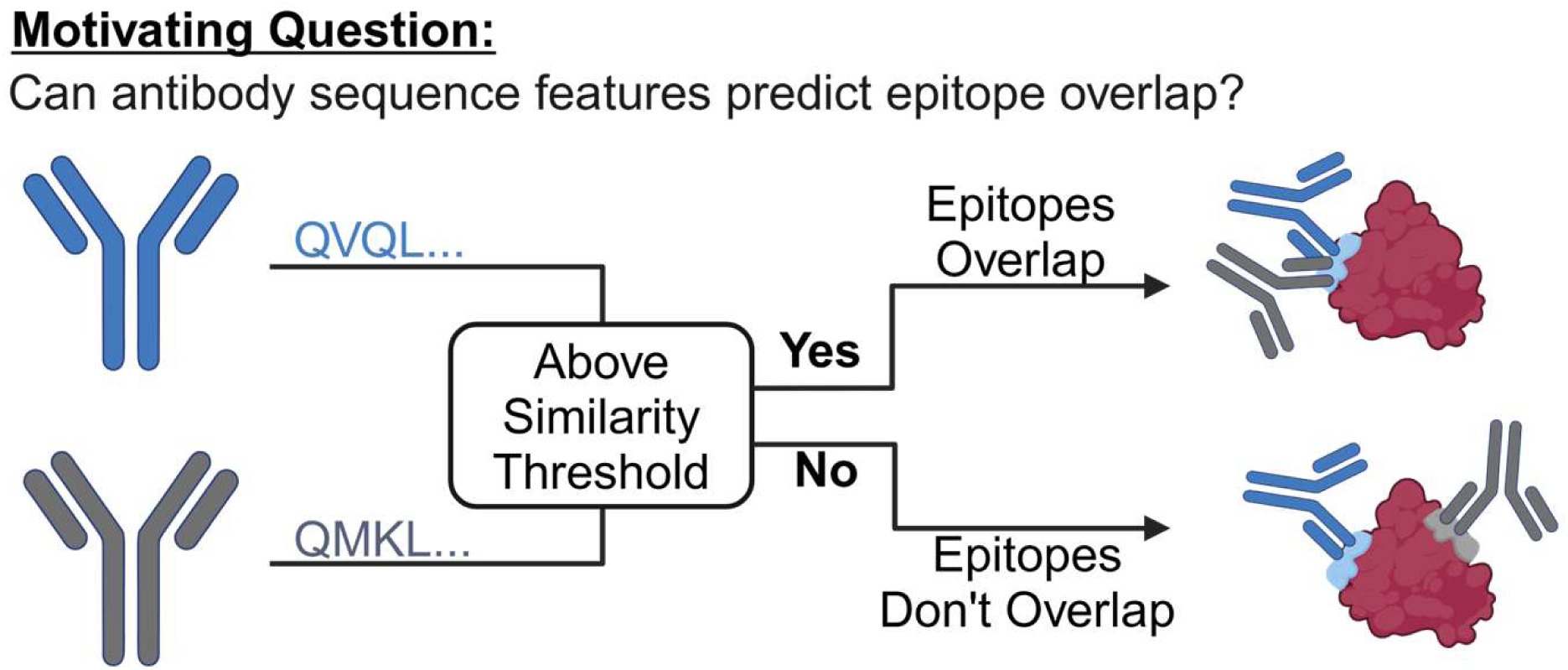
Motivating Question for this Work. Can antibody sequence features predict epitope overlap? If possible, then two antibodies (blue and gray) which have sequence feature similarities above a given similarity threshold are always known to target overlapping epitopes (top right). If their feature similarities are below this threshold they would be known to always target non-overlapping epitopes (bottom right). This study interrogates whether simple sequence features or more complicated features extracted from antibody amino acid sequences via machine learning are able to reliably distinguish overlapping epitope and non-overlapping epitope antibody pairs.

The emergence of antibody-specific language models, particularly AbLang, has opened new possibilities for computational antibody analysis ^22^. AbLang was trained on millions of naturally occurring antibodies through masked language modeling, where it learned to predict hidden amino acids based on surrounding sequence context ^23^. This training approach enabled the model to capture both evolutionary relationships and structural constraints within antibody sequences. However, like other current antibody language models, AbLang faces a critical limitation: its embeddings naturally cluster by sequence identity and germline gene usage, making it more adept at finding similar sequences than functionally similar antibodies with divergent sequences ^24,25^.

Recent advances in machine learning, particularly contrastive learning approaches, offer promising solutions to these limitations. Contrastive learning provides a framework for teaching models to recognize when two examples should be considered similar or different, even when observers see no clear patterns in their features. A useful analogy is image classification of flowers: while images of flowers from the same species may display distinct color, shape, and size differences, contrastive learning enables a model to recognize their fundamental similarities and distinguish them from similar species. By applying this approach to antibody analysis, we can explicitly train models to recognize structural or functional epitope similarity even when sequence similarity is low. Using carefully curated training data from structural databases and high-throughput epitope mapping experiments, we demonstrate how this approach can enrich antibody language model embeddings with epitope-specificity information while maintaining their broad understanding of antibody sequence space.

In this work, we address three key challenges in antibody epitope prediction. First, we establish reliable sequence-based thresholds for identifying overlapping-epitope antibodies, providing a simple yet powerful tool for repertoire analysis. Second, we develop and validate a model using the well-characterized SARS-CoV-2 receptor binding domain (RBD), where extensive epitope mapping data enables us to demonstrate how targeted training can overcome the germline bias of current language models. Finally, we present a generalized model capable of predicting epitope relationships across diverse protein families, which we validate through the successful identification of antibodies targeting overlapping epitopes with the HIV-1 broadly neutralizing antibody 8ANC195, a therapeutic candidate that targets a unique epitope on the HIV-1 envelope protein. These advances provide a comprehensive framework for computational epitope analysis, offering new possibilities for therapeutic antibody discovery and optimization.

## Results

### Sequence determinants for overlapping-epitope antibodies

To identify sequence features that reliably predict when antibodies target overlapping epitopes, we initially interrogated antibody sequence identity as a potential determinant. To that end, we focused on two key features of antibody recognition: variable (V) gene usage and CDR3 sequence similarity. We analyzed 1,909 non-redundant human antibodies from the Structural Antibody Database (SAbDab), generating approximately 1.8 million pairwise comparisons ^26,27^. These pairs were categorized based on both their V-gene sharing patterns and binding properties, specifically examining: (1) pairs binding overlapping epitopes, (2) pairs binding non-overlapping epitopes within the same protein family (Pfam), and (3) pairs binding different protein families (Fig. 2, S1A-B) ^28^.

**Figure 2.**
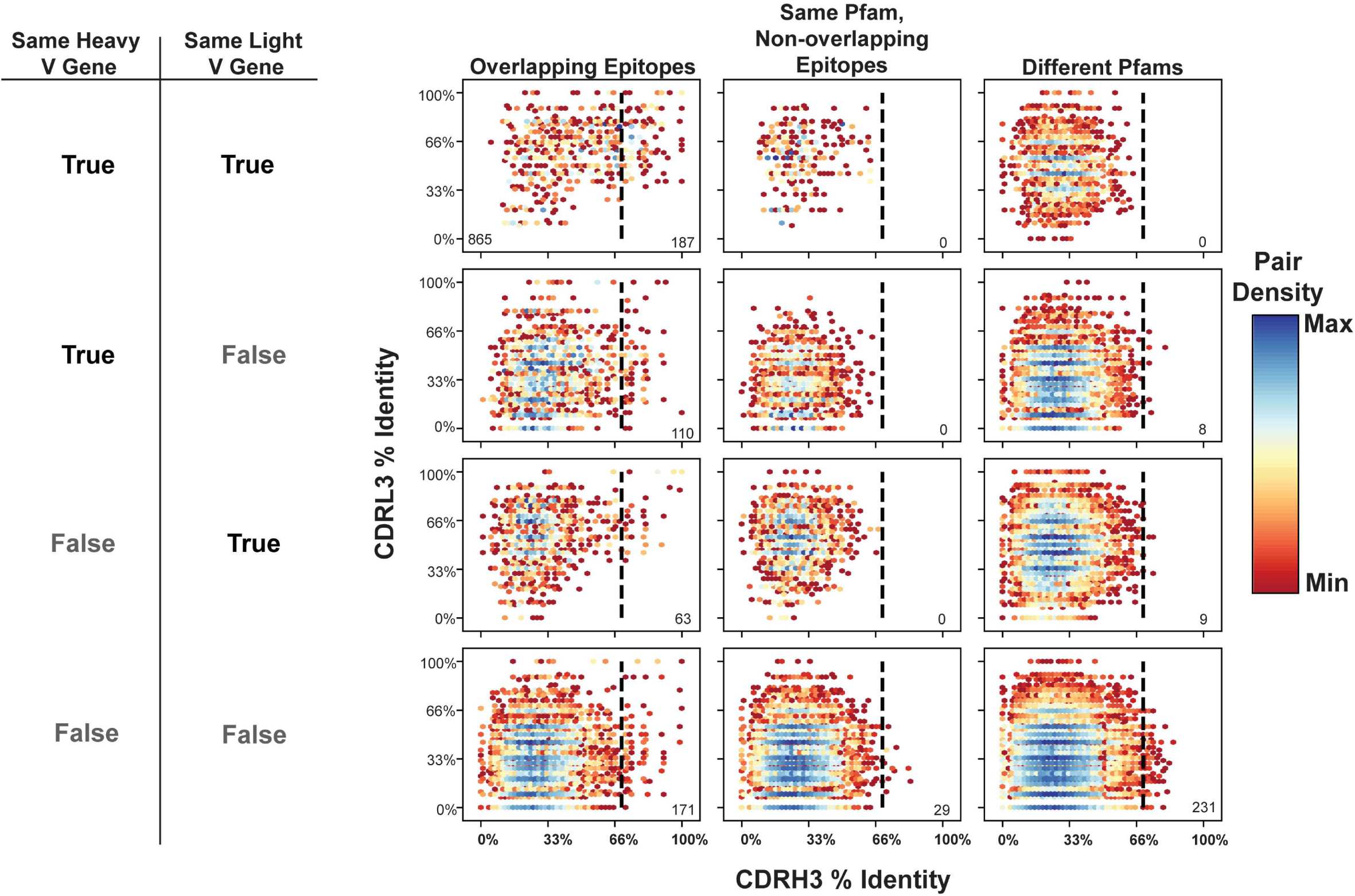
V-gene Usage and CDRH3 Sequence Identity Define Reliable Thresholds for Predicting Overlapping Epitopes. A comprehensive analysis of antibody sequence features predictive of epitope overlap within the Structural Antibody Database (SAbDab). Scatter plots show complementarity determining region 3 (CDR3) sequence identity relationships between antibody pairs (n = 1,909 antibodies, ∼1.8 million pairs) categorized by epitope relationship (columns) and V-gene sharing status (rows). The columns represent: overlapping epitopes (left), non-overlapping epitopes on the same protein family (middle), and different protein families (right). Rows indicate V-gene sharing patterns: both heavy and light V-genes shared (top), only heavy V-gene shared (second), only light V-gene shared (third), or neither V-gene shared (bottom). X-axis shows CDRH3 amino acid sequence identity; Y-axis shows CDRL3 amino acid sequence identity. Data density is represented by hexagonal binning with color scaling from minimum (dark red) through yellow to maximum density (dark blue). Dashed vertical lines indicate the 70% CDRH3 identity threshold. Numbers in bottom corners indicate pair counts within the half desigated by the line. When antibodies share at least one V-gene and target the same protein family, pairs exceeding 70% CDRH3 identity consistently bind overlapping epitopes (0 pairs in central columns). This relationship breaks down when no V-genes are shared (29 pairs in central column).

Our analysis revealed a hierarchical relationship between sequence features and epitope overlap. For antibody pairs sharing both heavy and light chain V-genes, we identified a heavy chain CDR3 (CDRH3) amino acid identity threshold of 70% that serves as a virtually perfect predictor - all pairs exceeding this threshold invariably bound overlapping epitopes within the same protein family (Fig. 2, top row). This predictive power persisted when antibodies shared only one V-gene, though with an important caveat (Fig. 2, rows 2 and 3): while pairs exceeding the CDRH3 threshold and targeting the same protein family consistently bound overlapping epitopes, 17 of 190 antibody pairs that exceeded this threshold bound antigens from entirely different protein families. This distinction suggests that when both the heavy and light chain germline V genes are shared, this provides additional constraints on antigen specificity beyond epitope recognition patterns.

While these sequence-based rules provide a clear framework for predicting epitope overlap, they also have two major limitations. First, the most predictive rule applies only to the small subset of antibody pairs sharing both V-genes. Second, even within this subset, the threshold fails to identify 82% of antibody pairs that do bind overlapping epitopes, resulting in a high false-negative rate. These limitations suggest that while sequence identity can provide absolute confidence in some cases, more sophisticated computational approaches may be needed for broader applicability in therapeutic antibody discovery.

To address these limitations, we next explored the ability of antibody large language models to learn the rules of epitope specificity. We focused on two domains: learning discrete epitope bins within one antigen and learning continuous epitope information across diverse protein families. These approaches, detailed in the following sections, demonstrate how modern computational methods can overcome the constraints of simple sequence-based rules ^34^.

### Contrastive Learning Enables Epitope-Specific Encoding of SARS-CoV-2 RBD Antibodies

While sequence-based thresholds provide reliable predictions in specific cases, their limited applicability motivated us to develop more sophisticated approaches for predicting epitope relationships. We leveraged the extensive epitope mapping data available for SARS-CoV-2 receptor binding domain (RBD) antibodies to develop and validate a contrastive learning framework that could encode epitope-specificity information directly into antibody sequence embeddings (Fig. 3) ^29–31^.

**Figure 3.**
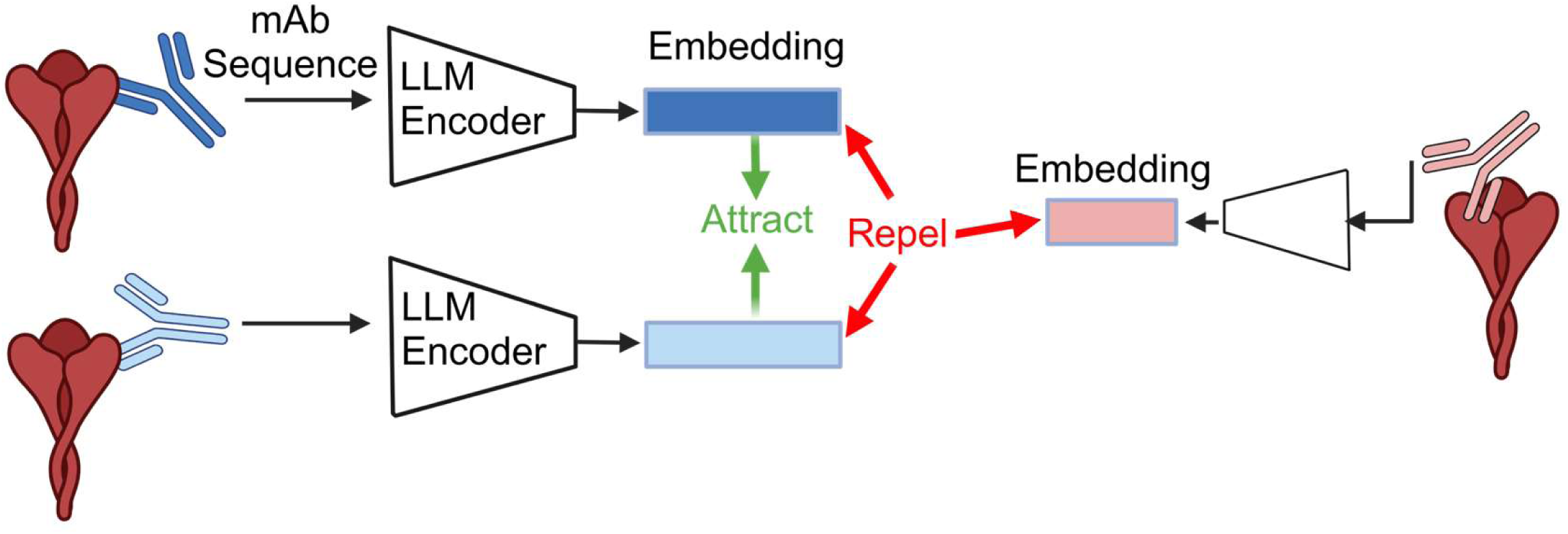
Contrastive Learning Framework for Encoding Epitope-Specificity Information in Antibody Sequence Embeddings. Schematic representation of the contrastive learning approach for training antibody language models to predict epitope overlap. Monoclonal antibody (mAb) amino acid sequences are processed through a large language model (LLM) encoder to generate sequence embeddings. During training, embeddings of antibodies binding overlapping epitopes (two blue antibodies) are pulled together (green arrows, “Attract”), while embeddings of antibodies binding non-overlapping epitopes (pink compared to blue) are pushed apart (red arrows, “Repel”). This framework enables the model to learn sequence features predictive of epitope overlap beyond simple sequence similarity metrics.

Our model, AbLang-RBD, builds upon the established AbLang heavy and light chain language models through targeted fine-tuning using a supervised contrastive learning framework ^22,32,33^. The architecture processes paired antibody sequences through a dual-stream transformer network - 12 separate transformer blocks per chain - followed by a six-layer multi-layer perceptron that generates unified sequence embeddings. We optimized these embeddings using a modified normalized temperature-scaled cross-entropy (NT-Xent) loss that simultaneously processes multiple positive examples, allowing the model to learn from groups of antibodies targeting the same epitope rather than individual pairs (Fig. 3). This approach differs from standard contrastive learning by concurrently attracting all antibodies sharing an epitope within a training batch while repelling those binding distinct epitopes, creating a more nuanced embedding space that captures epitope relationships ^33–35^. Critically, by training on same-epitope antibodies that fall outside our previously established V-gene and CDRH3 identity thresholds, the model learns new antibody sequence patterns indicative of shared epitope binding that are missed by our outlined V gene and CDRH3 thresholds.

We trained the model using a previously characterized set of 3,041 SARS-CoV-2 RBD antibodies binned into 12 epitopes based on deep mutational scanning results ^30,31^.

Notably, no antibody pairs in the training and test sets shared a heavy V gene and CDRH3 identity above 65%. Despite this, visualizing the cosine similarity distributions reveals that our contrastive learning approach effectively distinguishes epitope-bin information (Fig. 4A). The pretrained AbLang model showed poor separation between same-epitope and different-epitope pairs (56.0% balanced accuracy), whereas AbLang-RBD improved this distinction, achieving 74.4% balanced accuracy on test set comparisons and 82.7% when comparing test antibodies to the training set.

**Figure 4.**
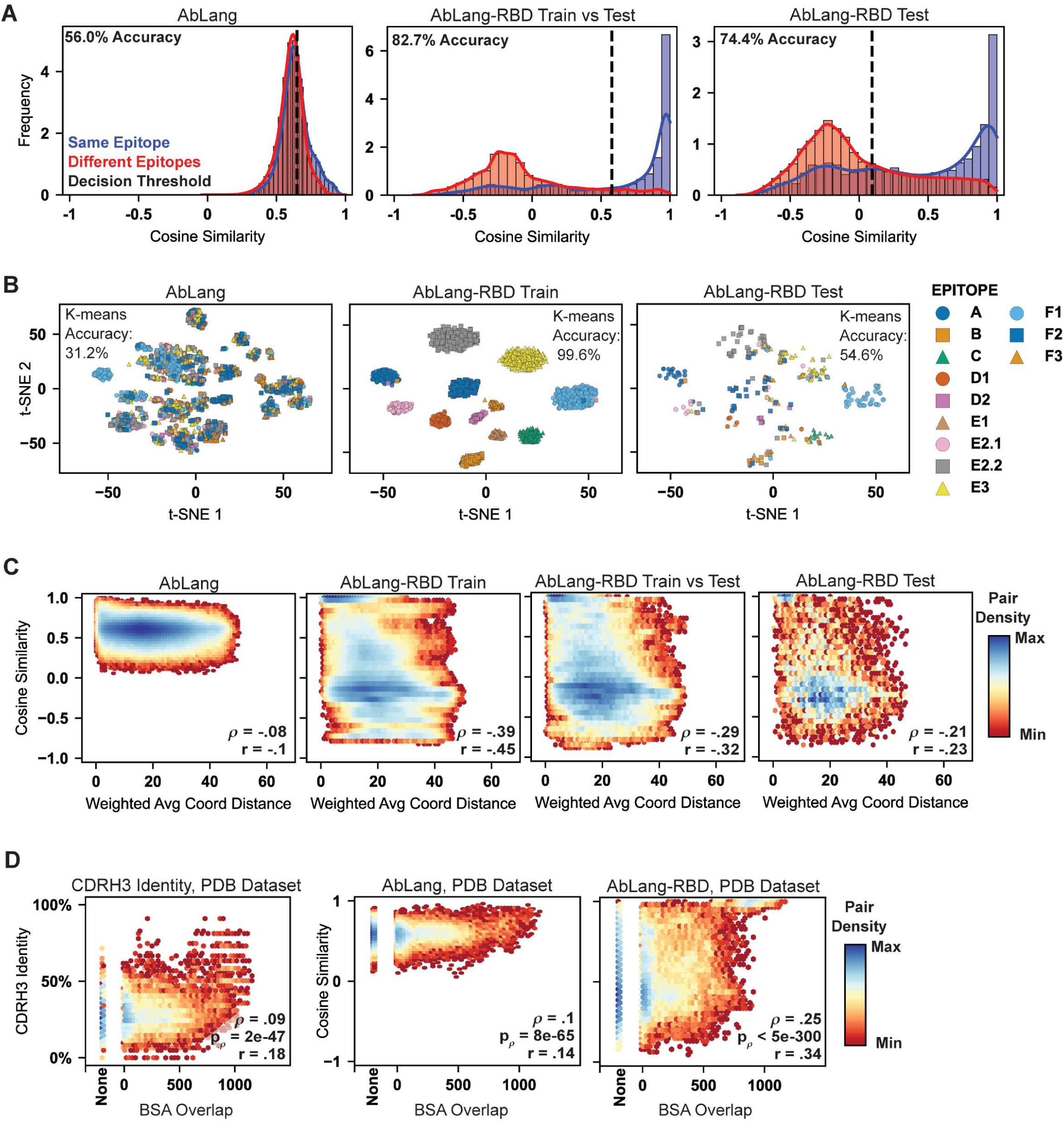
AbLang-RBD Learns to Predict Epitope Relationships from Binned Deep Mutational Scanning Data Fine-tuning performance of AbLang-RBD on SARS-CoV-2 RBD antibody epitope prediction. (A) Distribution of cosine similarities between antibody pairs binding same (blue) or different (red) epitopes. Left: Pretrained AbLang model shows poor separation (56.0% balanced accuracy). Middle: AbLang-RBD comparing training to test antibodies achieves 82.7% balanced accuracy. Right: AbLang-RBD comparing test antibodies achieves 74.4% balanced accuracy. Optimal decision thresholds (dashed lines) were determined using validation data. (B) t-SNE visualization of antibody embeddings colored by epitope class. Pretrained AbLang shows poor epitope clustering (31.2% k-means accuracy, left), while AbLang-RBD achieves near-perfect clustering on training data (99.6%, middle) and improved clustering on test data (54.6%, right). (C) Model performance assessed against deep mutational scanning (DMS) data. Scatter plots show relationship between antibody pair cosine similarities (y-axis) and weighted average spatial coordinates derived from DMS escape maps (x-axis). Hexagonal bins colored by pair density from minimum (dark red) to maximum (dark blue). Spearman’s (*ρ*) and Pearson’s (r) correlation coefficients shown. (D) Validation using structural data from PDB. Scatter plots compare CDRH3 sequence identity (left), pretrained AbLang (middle), and AbLang-RBD (right) against buried surface area (BSA) overlap between antibody pairs. AbLang-RBD shows improved correlation with structural epitope overlap (*ρ* = 0.25, p < 5e-300) compared to CDRH3 identity (*ρ* = 0.09, p = 2e-47) or pretrained AbLang (*ρ* = 0.1, p = 8e-65).

The effectiveness of our epitope-specific encoding is further demonstrated through dimensionality reduction analysis. T-distributed stochastic neighbor embedding (t-SNE) visualization reveals that while the pretrained model’s embeddings show minimal epitope-based clustering (31.2% k-means accuracy), AbLang-RBD achieves near-perfect clustering of training data (99.6%) and substantially improved clustering of test data (54.6%) (Fig. 4B) ^36,37^. Notably, when test antibodies were misclassified, 43% of errors still placed them within the correct RBD epitope class (out of 4 generally accepted classes), suggesting the model captures meaningful spatial relationships between epitopes despite this information not being provided in training ^38^.

To validate that our model learned genuine epitope-specificity information rather than arbitrary clustering, we evaluated its performance against two continuous data sources. First, we examined correlation with the underlying deep mutational scanning data by mapping escape scores onto the RBD structure to generate weighted average coordinates for each epitope (Fig. 4C, S1C). AbLang-RBD improved correlation with these spatial coordinates (Spearman’s *ρ* =-0.39, p < 5e-300) compared to the pretrained model (*ρ* =-0.08, p < 5e-300) for the training set as well as for the test set antibodies (*ρ* =-0.21, p = 5.2e-149). Second, we assessed performance on an independent set of 237 RBD-specific antibodies with structural epitope information from the PDB (Fig. 4D, S1A). AbLang-RBD demonstrated superior correlation with buried surface area overlap (*ρ* = 0.25, p < 5e-300) compared to both CDRH3 sequence identity (*ρ* = 0.09, p = 2e-47) and the pretrained model (*ρ* = 0.1, p = 8e-65).

These results demonstrate that supervised contrastive learning can effectively encode epitope-specificity information into antibody embeddings, establishing a powerful new framework for computational antibody analysis. While the model shows some limitation in distinguishing relative distances between non-overlapping epitopes, as evidenced by the presence of discrete bands in cosine similarity distributions rather than a continuous gradient (Fig. 4C), its ability to accurately identify antibodies targeting shared epitopes represents a significant advance over existing sequence-based methods. This capability, validated against both deep mutational scanning and structural data, provides a valuable new tool for therapeutic antibody discovery, particularly in cases where traditional sequence similarity metrics fail to identify functionally related antibodies. Most importantly, this framework establishes a foundation for developing even more sophisticated models that can capture the continuous nature of epitope relationships across diverse antigen families.

### Developing a Generalized Model for Continuous Epitope Overlap Prediction

To extend beyond predictions for a single antigen, we developed AbLang-PDB, a model capable of predicting the degree of epitope overlap for antibodies targeting antigens from diverse protein families represented in the Protein Data Bank (PDB) ^39–41^. Unlike AbLang-RBD’s discrete epitope binning approach, AbLang-PDB employs a regression framework to predict relative degrees of epitope overlap. We maintained the same dual-stream transformer architecture but modified the training objective to a mean squared error loss function to optimize for accurate comparisons between unseen antibodies and those in our curated training set.

The training data encompassed approximately 2,000 antibodies spanning 250 protein families, with relationships between antibody pairs encoded on a continuous scale.

Antibodies targeting different protein families received a label of-1, while those binding non-overlapping epitopes within the same protein family were assigned 0.2. For antibodies exhibiting epitope overlap, we assigned continuous labels from 0.5 to 1.0 based on their relative degree of structural overlap (Fig S1A-B). This nuanced labeling strategy enabled the model to learn the spectrum of possible epitope relationships rather than enforcing binary classifications.

The impact of this training approach is evident in the distribution of cosine similarities across different antibody pair categories (Fig. 5A). While the pretrained AbLang model showed minimal separation between the three categories (39.1% balanced accuracy), AbLang-PDB achieved clear differentiation (62.5% balanced accuracy), with overlapping-epitope pairs predominantly exhibiting cosine similarities above 0.75. More importantly, the model demonstrated strong correlation with ground truth labels (*ρ* = 0.304 out of a possible 0.525 due to ties) across all comparisons, suggesting it learned meaningful relationships between sequence features and epitope overlap (Fig. 5B).

**Figure 5.**
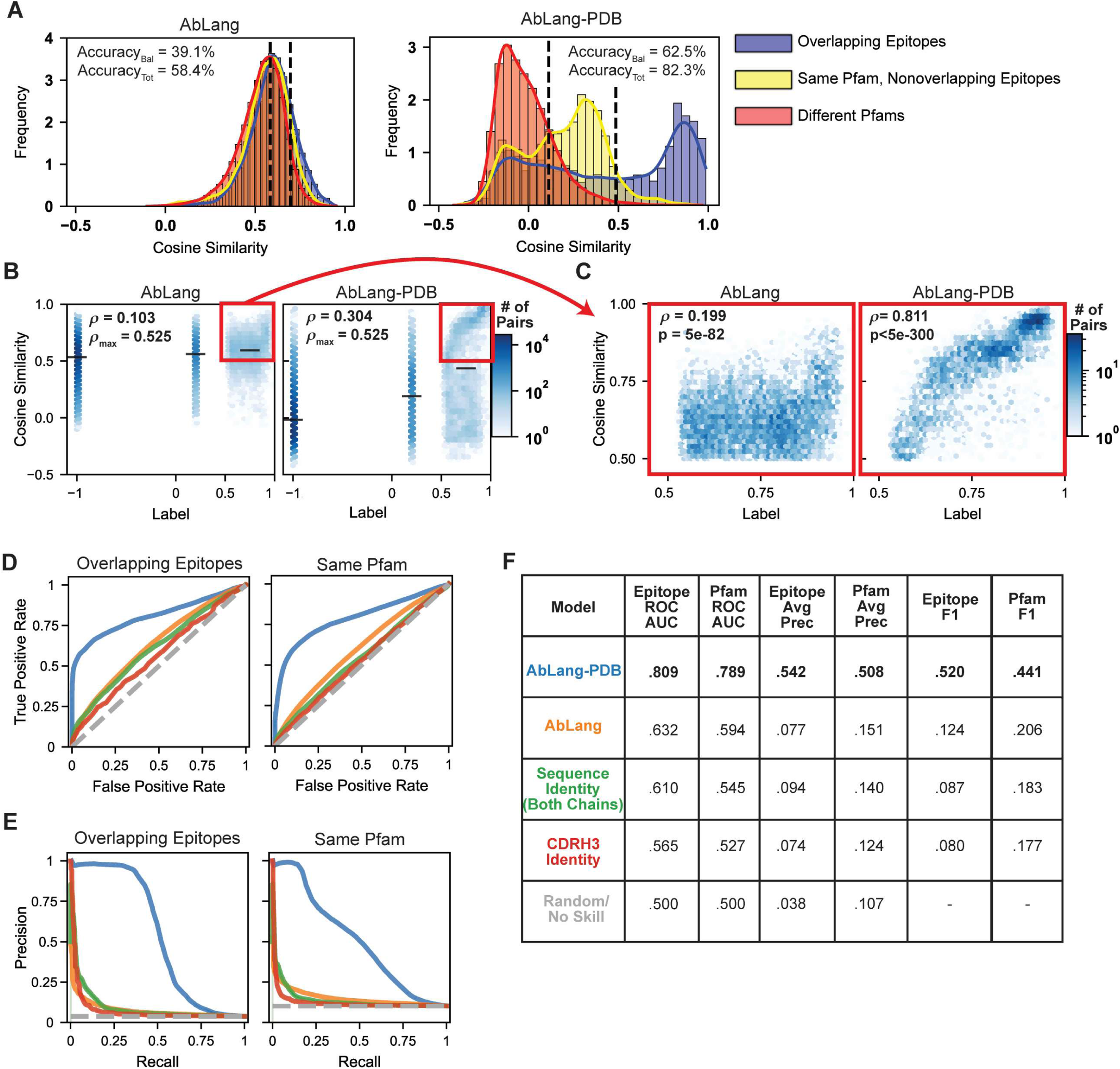
AbLang-PDB Enables Accurate Prediction of Epitope Relationships Across Diverse Protein Families. Evaluation of AbLang-PDB’s performance on the Structural Antibody Database (SAbDab). (A) Distribution of cosine similarities between antibody pairs categorized as overlapping epitopes (blue), non-overlapping epitopes on the same protein family (yellow), or different protein families (red). Left: Pretrained AbLang shows poor separation (balanced accuracy 39.1%, total accuracy 58.4%). Right: AbLang-PDB achieves improved separation (balanced accuracy 62.5%, total accuracy 82.3%). Optimal classification thresholds for balanced accuracy on the validation set shown as dashed lines. (B) Relationship between model-predicted cosine similarities and ground truth labels. Hexagonal bins colored by pair density (white to dark blue). Black bars indicate mean ± 95% confidence intervals. Spearman correlations (*ρ*) and maximum possible correlations (ρ_max_) shown. (C) Detailed view of high-confidence predictions (cosine similarity and label ≥ 0.5) showing strong correlation for overlapping-epitope pairs (*ρ* = 0.811, p < 5e-300). (D) Receiver operating characteristic curves comparing classification performance of AbLang-PDB (blue) against pretrained AbLang (orange), full-sequence identity (green), CDRH3 identity (red), and random prediction (gray) for overlapping epitopes (left) and shared protein family (right). (E) Corresponding precision-recall curves demonstrating AbLang-PDB’s several fold improvement relative to other prediction methods. (F) Summary metrics including area under ROC curve (ROC AUC), average precision (Avg Prec), and F1 scores for epitope and protein family predictions across methods.

Notably, the model performs exceptionally well on high-confidence predictions among overlapping-epitope pairs. When considering only these pairs with model-predicted cosine similarities above 0.5, the correlation with actual epitope overlap increases dramatically to *ρ* = 0.811 (p < 5e-300) (Fig. 5C). This indicates that while the model may not perfectly separate all antibody pairs, it can be used within this application for highly reliable predictions on the extent of epitope overlap.

Comprehensive evaluation through receiver operating characteristic and precision-recall analyses revealed substantial improvements over existing methods (Fig. 5D-F). For overlapping-epitope classification, AbLang-PDB achieved an ROC-AUC of 0.809, an average precision of 0.542 and an F1-score of 0.52, significantly outperforming both the pretrained model (0.632, 0.077, 0.124) and sequence-identity-based predictions (0.610, 0.094, and 0.087). Similar improvements were observed for Pfam prediction, with 1.32x, 3.36x, and 2.14x enhancements over the best alternative classifier in the respective categories (Fig. 5D-F).

These results demonstrate that our continuous learning approach successfully captures epitope relationships across diverse protein families while maintaining high precision for overlapping-epitope predictions. The model’s ability to provide reliable confidence scores through cosine similarities makes it particularly valuable for therapeutic antibody discovery, where false positives can be costly. In the following section, we validate this capability through the successful identification of novel antibodies targeting a therapeutically relevant HIV-1 epitope.

### Experimental validation of AbLang-PDB through epitope-targeted HIV-1 antibody discovery

To validate AbLang-PDB’s practical utility, we applied it to identify antibodies sharing epitope overlap with the HIV-1 broadly neutralizing antibody 8ANC195 ^42–44^. This antibody represents an ideal test case due to its unique binding site at the gp120-gp41 interface and its therapeutic potential, despite having somewhat limited neutralization breadth compared to other broadly neutralizing antibodies ^45^. We analyzed a dataset of 7,056 class-switched antibodies from persons living with HIV-1, computing cosine similarities between each antibody and 8ANC195’s embedding (Fig. 6A-B) ^46,47^. These antibodies were identified through LIBRA-seq (linking B cell receptor to antigen specificity through sequencing), a high-throughput technology that enables simultaneous identification of antigen specificity and BCR sequences at single-cell resolution. In this approach, B cells are exposed to oligonucleotide-barcoded antigens, allowing quantitative assessment of antigen binding through unique molecular identifiers during subsequent single-cell sequencing. The dataset comprised 21 LIBRA-seq experiments where antigen-specific B cells were isolated from peripheral blood mononuclear cells (PBMCs) using fluorescence-activated cell sorting (FACS). To ensure specificity, each B cell’s antigen-binding profile was determined using both target antigens and control antigens (both positive and negative) conjugated to the same fluorophore. While this experimental design enriched for HIV-1 specific antibodies in the dataset, the majority of sequences are not expected to be HIV-1 specific. From this analysis, we identified 20 candidates with the highest cosine similarities (range: 0.567-0.655). After eliminating eight antibodies with high sequence identity to other selected candidates and two showing potential reactivity to respiratory syncytial virus or hepatitis C virus in their LIBRA-seq profiles, we prioritized 10 antibodies for experimental characterization. Notably, the intermediate cosine similarity scores suggested partial rather than complete epitope overlap, consistent with 8ANC195’s unique epitope characteristics (Fig. 6A).

**Figure 6.**
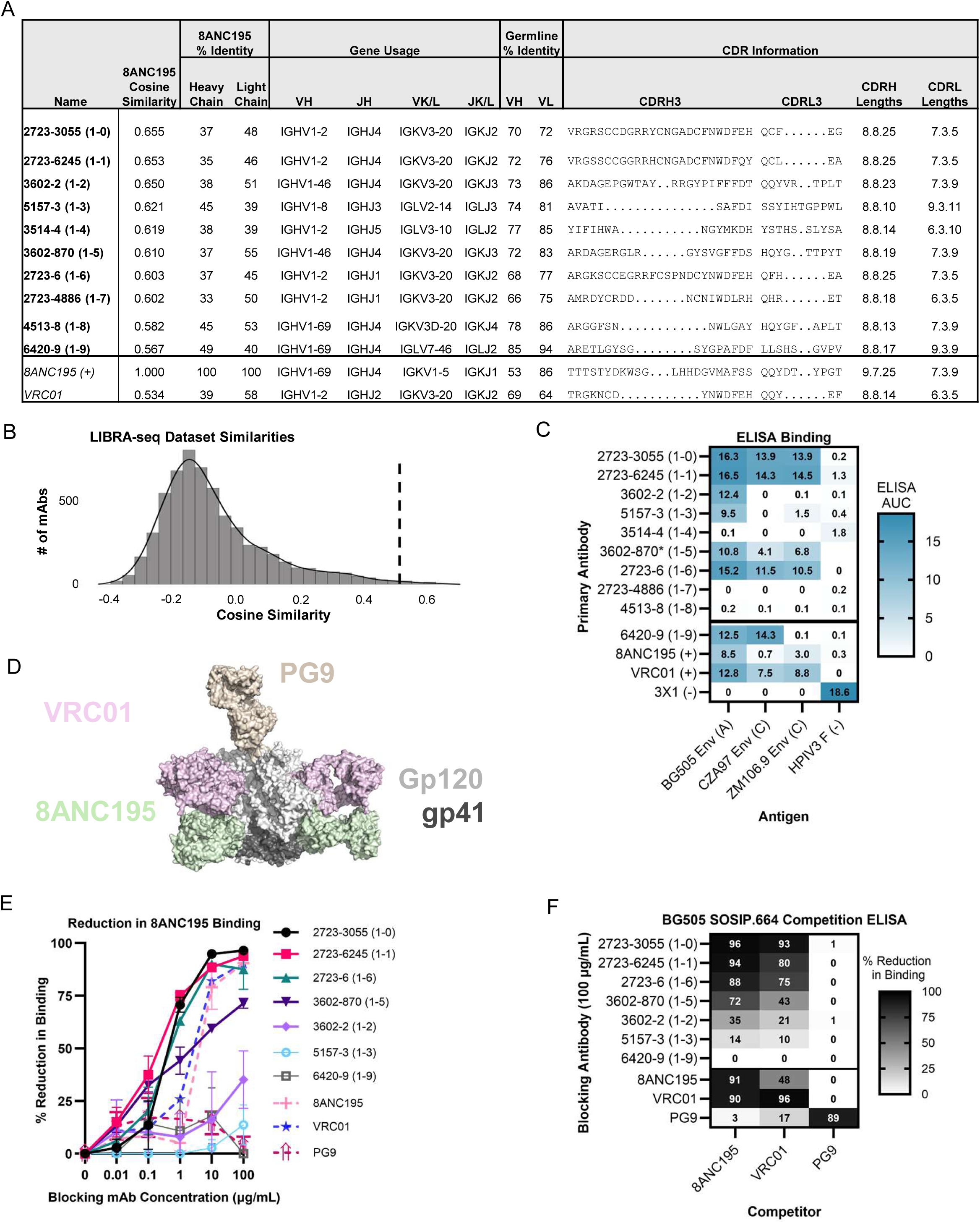
AbLang-PDB Successfully Identifies HIV Antibodies which compete for binding with 8ANC195. Experimental validation of AbLang-PDB predictions using HIV broadly neutralizing antibody 8ANC195. (A) Sequence characteristics of top candidate antibodies selected by cosine similarity to 8ANC195. Table shows model predictions, sequence identity metrics, gene usage, and CDR information for each antibody. Reference antibodies 8ANC195 and VRC01 included for comparison. (B) Distribution of cosine similarities across the complete LIBRA-seq dataset (n = 7,056 antibodies), with dashed line indicating recommended threshold (0.5) for mining overlapping-epitope candidates. (C) ELISA binding profiles against HIV-1 envelope SOSIP.664 constructs (BG505, CZA97, ZM106.9) and HPIV3 F control protein. Binding strength indicated by area under the curve (white to blue). 3X1 antibody included as HPIV3-specific control. (D) Structural representation of HIV-1 BG505 envelope showing competitor antibody epitopes: 8ANC195 (green, gp120-gp41 interface), VRC01 (pink, CD4-binding site), and PG9 (tan, V1-V2 region) from PDB IDs 5VJ6 and 8VGW. Envelope surface shown with gp120 (light gray) and gp41 (black). (E) Competition ELISA curves showing percent reduction in binding of biotinylated 8ANC195 (10 μg/mL) to BG505 SOSIP.664 in presence of increasing concentrations of blocking antibodies. Filled symbols indicate mAbs displaying competition with 8ANC195. (F) Competition matrix showing percent reduction in binding at fixed concentrations (blocking: 100 μg/mL; detection: 8ANC195 10 μg/mL, VRC01 and PG9 1 μg/mL). Values range from no competition (white) to complete competition (black).

We first assessed HIV-1 specificity through ELISA against three diverse HIV-1 envelope SOSIP.664 constructs: BG505 (Clade A), CZA97 (Clade C), and ZM106.9 (Clade C), using human parainfluenza virus 3 fusion (HPIV3 F) protein as a negative control (Fig. 6C, S2A-B ^46^). Seven of the ten selected antibodies (70%) demonstrated HIV-1 envelope specificity, with six (60%) showing cross-clade binding. Out of our unbiased selection, two had previously been characterized—2723-3055 and 3602-870—both of which had been shown to potently neutralize a broad panel of tier 2 HIV-1 viruses (12/12 and 11/14 viruses tested) ^46^.

To assess epitope overlap, we performed competition ELISAs against BG505 SOSIP.664 using a panel of well-characterized HIV-1 antibodies targeting distinct epitopes: 8ANC195 (gp120-gp41 interface), VRC01 (CD4 binding site), and PG9 (V1-V2 region) (Fig. 6D-F, S2C) ^48,49^. Five antibodies (50%) showed competition with 8ANC195, defined as >30% reduction in binding, with four of those displaying strong competition (>70% reduction). These same antibodies also competed with VRC01 at a slightly weaker level but not with PG9, consistent with the structural overlap between the 8ANC195 and VRC01 epitopes.

The success rate of our computational predictions - 50% for identifying 8ANC195-competing antibodies and 70% for HIV-1 Env specificity - highlights AbLang-PDB’s potential to streamline therapeutic antibody discovery by accurately identifying functionally relevant candidates that conventional sequence similarity metrics can miss. This is particularly noteworthy given that the model identified two previously validated broadly neutralizing antibodies without any prior knowledge of their functional properties. These results support AbLang-PDB’s utility for therapeutic antibody discovery, especially in cases where conventional sequence similarity metrics would fail to identify functionally related candidates.

## Discussion

The identification of antibodies targeting overlapping epitopes remains a critical challenge in therapeutic antibody development. Current approaches typically rely on experimental screening or database searches for sequence-similar antibodies, both of which have significant limitations ^7,50^. Our work establishes three complementary computational strategies that address this challenge at different levels of complexity and applicability.

First, we provide rigorously validated sequence identity-based thresholds for predicting epitope overlap. When antibody pairs share both heavy and light chain V-genes and have CDRH3 amino acid identity exceeding 70%, they not only consistently bind overlapping epitopes but are also guaranteed to target the same protein family. This constraint relaxes slightly when only one V-gene is shared - while the 70% CDRH3 identity threshold still perfectly predicts overlapping epitopes when antibodies target the same protein family, some pairs meeting this criterion may bind antigens from different protein families altogether. While these findings appear straightforward, they represent the first systematic validation of such thresholds across more than 200 antigen specificities. These simple yet powerful criteria provide immunologists with reliable tools for initial repertoire analysis, particularly valuable when analyzing vaccine responses or comparing antibody lineages across individuals.

Second, through AbLang-RBD, we demonstrate how supervised contrastive learning can enhance language model embeddings with epitope-specificity information. Using the well-characterized SARS-CoV-2 RBD as a model antigen, we showed that this approach achieved 74.4% accuracy in predicting epitope relationships between previously unseen antibodies. The model’s ability to generalize beyond its training data is evidenced by its improved correlation with both deep mutational scanning data and structural epitope measurements compared to sequence-based metrics or the pretrained model. This indicates that our contrastive learning framework indeed captures genuine epitope-specificity information rather than merely clustering similar sequences.

Third, we developed AbLang-PDB, which extends epitope prediction capabilities across diverse protein families while capturing continuous relationships between epitopes. This model demonstrates substantial improvements over existing methods, achieving a five-fold increase in average precision for overlapping-epitope prediction while simultaneously improving antigen protein family average precision 3.4-fold. Particularly noteworthy is AbLang-PDB’s ability to provide reliable confidence scores through cosine similarities for overlapping epitope antibodies, with high-confidence predictions (cosine similarity > 0.5) showing strong correlation (*ρ* = 0.811) with actual epitope overlap.

The practical utility of these approaches is demonstrated by our successful identification of HIV-specific antibodies sharing epitope overlap with 8ANC195. Among our computationally selected candidates, 70% showed HIV specificity and 50% competed with 8ANC195 for binding. Additionally, despite this dataset containing only two previously characterized broadly neutralizing antibodies, both antibodies were among the model’s top 10 cosine similarity scores validating the model’s ability to mine datasets for therapeutically relevant antibodies without requiring the complex experimental screening methods typically needed for identifying such candidates.

We also note, however, that our approaches have important limitations. The sequence identity-based thresholds, while providing perfect precision for predicting overlapping epitopes, exhibit low recall - failing to identify 82% of antibody pairs that do share epitope overlap. AbLang-RBD demonstrates high performance on SARS-CoV-2 index strain RBD epitope prediction but faces two key constraints: it is not clear that it will generalize to RBD-specific antibodies incapable of binding the index strain, and its training approach using deep mutational scanning data has not been validated for the more commonly available antibody-antibody competition binding data. AbLang-PDB’s training paradigm is optimized for comparing novel antibodies against those in its training set rather than directly comparing two previously unseen antibodies, making it more suitable for analyses that leverage known reference antibodies to assess similarity and epitope overlap. While we validated the model using HIV-1 envelope as a test case, this antigen is well-represented in our training data, though notably, our validation epitope (8ANC195’s epitope) had minimal representation.

Despite these limitations, our work provides a comprehensive framework for computational epitope analysis that will be of significance for the field of therapeutic antibody discovery. The combination of simple sequence rules and sophisticated machine learning models offers researchers a tiered approach to identifying overlapping-epitope antibodies, from rapid initial screening to detailed prediction of epitope relationships. Looking forward, these methods can be further enhanced through integration with emerging structural prediction tools and expanded training datasets, potentially enabling even more accurate prediction of antibody-antigen interactions.

## Supplemental Figure Captions

**Figure S1. Quantitative Framework for Defining Epitope Relationships and Dataset Characteristics.** (A) Systematic approach for determining epitope overlap and machine learning labels for antibody pairs targeting antigens within the same Pfam family. Two complementary metrics are employed: (1) SASA-based residue-level epitope overlap, calculated as the overlapping solvent-accessible surface area buried by the antibodies (threshold >20 Å²), and (2) distance-based atom-level epitope overlap, defined by antigen atoms within 4.5 Å of each antibody (threshold >5 Å²). Pairs exceeding both thresholds receive a machine learning label of MAX(1, 0.5 + Average(RELATIVE_BSA_OVERLAP, RELATIVE_ATOM_OVERLAP)^0.75^) otherwise, they are assigned a non-overlapping label of 0.2. (B) Relationship between machine learning labels and buried surface area (BSA) overlap visualized through hexagonal binning density plot (r = 0.89, Y_Fit_ = 4.07e-4*x + 0.584 for pairs with labels ≥ 0.5). Color intensity indicates pair density from minimum (red) to maximum (blue). (C) Three-dimensional visualization of deep mutational scanning (DMS) weighted average coordinates, with points colored by epitope classification. Coordinate calculation methodology detailed in Methods. Data demonstrates points maintain clear spatial segregation of epitope classes.

**Figure S2: ELISA data for 8ANC195-targeted antibody discovery campaign. (A-B) ELISA curves pertaining to** Figure 6C **AUC values.** Binding to trimeric HIV-1 envelope (BG505, CZA97, ZM106.9) or to a negative control antigen, human parainfluenza virus 3 fusion protein. (B) ELISA curves for antibody 3602-870, reproduced from data collected from 3602-870’s initial publication. Antigens listed without underscores refer to the use of 3602-870 as the primary antibody whereas antibodies after underscores are positive controls for that antigen. H1 NC99 stands for the Influenza A hemagglutinin from the strain A/New Caledonia/20/99 (H1). (C) Competition ELISA curves showing detection of biotinylated antibody binding to BG505 SOSIP.664 with the biotinylated antibody as the title (8ANC195, VRC01, or PG9).

Absorbance at 450 nm shown in presence of increasing concentrations of competitor antibodies. Filled symbols indicate mAbs displaying competition with 8ANC195.

## Resource availability

### Lead Contact

Further information and requests for resources and reagents should be directed to and will be fulfilled by the lead contact, Ivelin S. Georgiev.

### Materials availability

Materials will be made available upon request under a completed material transfer agreement (MTA).

### Data and code availability

Sequences for antibodies identified and characterized in this study will be deposited to GenBank following journal acceptance and prior to publication.

Associated code for AbLang-RBD and AbLang-PDB will be made available at https://github.com/IGlab-VUMC/AbLangRBD1 and https://github.com/IGlab-VUMC/AbLangPDB1 and the links for downloading model weights will also be made available there prior to peer-reviewed publication.

Any additional data or code reported in this paper will be shared by the lead contact upon request.

## Acknowledgements

We thank Perry Wasdin for his help in curating antigen-specific training sets which helped interrogate the efficacy of different models and loss functions and Alexandra Abu-Shmais for her insights on drawing conclusions from pairwise antibody comparisons. We additionally thank Andrea Shiakolas, Ian Setliff, Kelsey Pilewski, Rohit Venkat, and Lauren Walker for LIBRA-seq data. This research was funded, in part, by the Advanced Research Projects Agency for Health (ARPA-H), NIH R01AI175245, and G. Harold and Leila Y. Mathers Charitable Foundation (MF-2107-01851). The funders had no role in the conceptualization or execution of any studies or drafting of the manuscript. The views and conclusions contained in this document are those of the authors and should not be interpreted as representing the official policies, either expressed or implied, of the U.S. Government.

## Methods

### Data curation

We curated a comprehensive dataset from the Structural Antibody Database (SAbDab, February 19, 2024 cutoff date) for training and validating the AbLang-PDB model ^26,27^. Starting with 16,105 antibody-antigen complexes, we applied the following filtering criteria: resolution ≤ 4.5 Å, human antibodies with both chains present, and ≥5 amino acid differences between antibodies. This yielded 1,909 non-redundant complexes, of which 184 had no same-Pfam pairs and 485 had no overlapping-epitope pairs.

Antigen classification utilized pfam_scan software to group antigens by domain architecture using hidden Markov models ^28,51^. Multiple Pfam assignments were consolidated such that any shared Pfam between antigens classified their respective antibodies as targeting the “same Pfam” and thus a machine learning label of 0.2. When no overlap was present these pairs were assigned a machine learning label of-1. For quantifying epitope overlap, we employed two complementary approaches. First, we calculated buried surface area (BSA) per residue using DSSP by comparing the amount of surface area at each residue for the antigen either in complex with the antibody or without the antibody ^52^. We did this after aligning antigen sequences using the BLOSUM62 matrix and Needleman-Wunsch algorithm ^53,54^. Per-residue BSA overlap was calculated as MIN(BSA_res1, complex1_, BSA_res1, complex2_). Antibody pairs with total BSA overlap summed over all residues ≤ 20 Å² were labeled as non-overlapping (Fig S1A, label = 0.2). Second, we defined epitopes as antigen heavy atoms within 4.5 Å of antibody atoms and calculated overlap volume using PyMOL’s overlap function, with pairs showing overlap ≤ 5 Å³ labeled as non-overlapping (label = 0.2) ^55^.

For overlapping epitopes, final labels were assigned on a continuous scale from 0.5 to 1.0 using the formula: Label = max(1, 0.5 + (rBSA_OVERLAP + rATOM_OVERLAP)×0.75), where rBSA_OVERLAP and rATOM_OVERLAP represent overlap relative to the smaller of the two self-overlap values seen for each epitope pair. For partitioning antibodies between datasets, antibodies sharing both heavy and light V-genes and CDRH3 amino acid identity >65% were assigned to the same clone group. These groups were then distributed across training (80%), validation (10%), and test (10%) sets, ensuring no clone group appeared split between multiple sets. Additionally, pairs with >92.5% sequence identity in either chain were excluded to maintain diversity.

For AbLang-RBD, we utilized published deep mutational scanning data comprising 3,195 antibodies from 2 papers, of which only the 3,093 which demonstrated binding to SARS-CoV-2 index strain were kept ^30,31^. These antibodies were clustered based on heavy chain V-gene usage and CDRH3 amino acid identity >70%, with clusters distributed across training (80%), validation (10%), and test (10%) sets such that no antibodies in the same cluster existed in the training and test sets. A separate test set was curated from the PDB by selecting RBD-specific antibodies from CoV-AbDab that demonstrated index strain binding, and were unique from those in the deep mutational scanning dataset ^29^. This left 237 antibodies and 27,345.

### Model Architecture

Our approach built upon the pretrained AbLang framework, which comprises separate heavy and light chain transformer models for antibody sequence analysis. We utilized the published AbLang model weights from Huggingface (qilowoq/AbLang_heavy and qilowoq/AbLang_light) accessed through the transformers library (AutoModel, AutoTokenizer) ^22,56^. The base architecture follows RoBERTa with modifications for antibody sequence processing: each chain is processed through 12 transformer blocks containing 12 attention heads, with hidden dimension 768 and intermediate dimension 3072. A learned positional embedding layer handles sequences up to length 160 ^57^.

For sequence processing, antibody amino acid sequences were first tokenized using the transformers module. Heavy and light chain sequences were processed independently through their respective models to generate chain-specific embeddings. For each chain, the final hidden layer outputs (768-dimensional vectors) from all non-masked positions were mean-pooled. While for the pretrained AbLang model we simply concatenated these chain embeddings, the architecture for AbLang-RBD and AbLang-PDB introduces additional processing layers to enable cross-chain information flow.

Specifically, the concatenated 1536-dimensional vector (768 dimensions per chain) is processed through a 6-layer multi-layer perceptron with ReLU activation between layers, except for the final layer. The normalized output of this network serves as the unified antibody embedding.

To enable efficient fine-tuning while preserving pretrained weights, we employed QLORA (Quantized Low-Rank Adaptation) with rank R=16, alpha=32, and dropout=0.3^32^. This dual-stream architecture - with 12 transformer blocks per chain followed by the cross-chain mixing network - allows the model to capture both chain-specific features and relationships between heavy and light chain sequences.

### AbLang-RBD training

The AbLang-RBD model was trained using a supervised contrastive learning approach with a modified NT-Xent (InfoNCE) loss function ^33,35^. During training, we froze all pretrained weights except for the QLORA adaptation parameters and the six “mixing” layers that enable cross-talk between heavy and light chain embeddings. Optimization was performed using the AdamW optimizer with a learning rate of 1e-5 and batch size of 256.

Specifically, the loss function is a multi-positive variant of the NT-Xent (InfoNCE) loss, where each sample *z_i_*, may have multiple positive samples {*z_j_*} sharing the same label and sim stands for cosine similarity ^33,58,59^. For each positive pair, we take

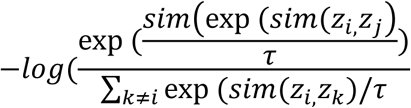

and then average the resulting losses over all positive pairs in the batch using a temperature τ = 0.5 and a batch size of 256.

Training proceeded for 400 epochs on a single NVIDIA A6000 GPU, requiring approximately 5 hours including inter-epoch evaluations. Model selection was based on ROC-AUC performance on the validation set (weighted by epitope class size), with the epoch 280 checkpoint achieving optimal performance.

### Histogram generation and pairwise accuracy or F1 calculation

Distributions of antibody pair relationships were visualized and analyzed using histograms implemented in Python 3.8.18 with seaborn 0.13.1. All histograms were generated using probability density normalization with 30 uniform-width bins.

Classification thresholds were determined differently for AbLang-RBD and AbLang-PDB versus the pretrained model. For AbLang-RBD and AbLang-PDB, thresholds were optimized to maximize balanced accuracy on the validation dataset. The pretrained model threshold in Figure 5A was similarly optimized using train versus validation parameterization, while in Figure 4A it was optimized for maximal balanced accuracy across the complete dataset (note: this approach overestimates model performance).

For three-category classification, optimal decision boundaries were determined via grid search across 90,000 threshold combinations (300 x 300 cosine similarity values). The threshold pair yielding maximum balanced accuracy across all three categories (overlapping epitopes, non-overlapping epitopes within the same Pfam, and different Pfams) was selected. Balanced accuracy was calculated as the mean of individual category accuracies, in contrast to total accuracy which can be biased by class imbalance.

### Calculation of a representative deep mutational scanning coordinate

Deep mutational scanning (DMS) escape data were obtained for 1,375 SARS-CoV-2 RBD antibodies from the publicly available Bloom laboratory database ^30,31,60,61^. For each antibody, a representative three-dimensional coordinate was calculated using position-specific escape scores as weights. Specifically, escape scores were first aggregated by residue position, with the weighted average coordinate (*x4*) calculated as:

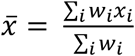

where *w_i_*is the sum of the escape scores for all mutations at residue i, and *x_i_*is the 3D coordinates of the alpha carbon for residue i coming from the SARS-CoV-2 RBD structure (PDB ID 8SGU)^62^. Pairwise distances between antibody coordinates were calculated using Euclidean distance metrics. To validate this coordinate representation scheme, we visualized the three-dimensional distribution of antibody positions colored by epitope class, confirming minimal distortion of known epitope relationships (Fig. S1C).

### Regression analysis

Statistical analyses were performed using SciPy (version 1.10.1) for correlation calculations and significance testing ^63^. Spearman’s rank correlation (spearmanr) and Pearson correlation (pearsonr) coefficients were calculated for various pairwise comparisons. In Figure S1B, the relationship between buried surface area (BSA) and training labels was fit using linear regression (scipy.stats.linregress), excluding pairs with labels below 0.5. For correlation analyses in Figures 4D and 5B-C, Spearman correlations were calculated with associated p-values; p-values below the numerical precision limit of 64-bit floating point numbers are reported as p < 5e-300.

For Figure 5B, we calculated the maximum achievable Spearman correlation (ρmax) by considering the optimal ranking scenario where: (1) all antibody pairs with label-1 rank below those with label 0.2, (2) all pairs with label 0.2 rank below those with labels ≥ 0.5, and (3) pairs with labels between 0.5 and 1.0 are perfectly rank-ordered. Mean values with 95% confidence intervals were calculated for discrete label categories (-1 and 0.2) and for the continuous range of labels ≥ 0.5 (plotted at x = 0.75). For Figure 5C, analysis was restricted to pairs with both predicted cosine similarities and ground truth labels between 0.5 and 1.0 to assess performance on high-confidence predictions.

### T-SNE analysis and K-means accuracy calculation

Dimensionality reduction and clustering analyses were performed using scikit-learn (version 1.3.2). For t-SNE visualization, 1536-dimensional antibody embeddings were reduced to two dimensions using the following parameters: PCA initialization, automatic learning rate determination, perplexity of 30, and maximum 1000 iterations with a learning rate of 1000. For AbLang analysis, the complete dataset was visualized in aggregate. For AbLang-RBD, while dimensionality reduction was performed on the complete dataset, training and test sets were subsequently visualized separately to assess generalization performance.

Clustering analysis was performed using k-means with cosine similarity as the distance metric. The algorithm was initialized with 12 clusters using the k-means++ strategy for greedy centroid initialization and allowed to run for a maximum of 300 iterations.

Clustering accuracy was assessed by assigning the most highly represented epitope class within each cluster as the cluster’s representative epitope. Antibodies within each cluster were considered accurately clustered if they matched this epitope and incorrectly clustered otherwise. This approach, while disadvantaging underrepresented epitopes due to class imbalance, provides a conservative estimate of clustering performance.

For visualization clarity, we cycled through three marker shapes (circles, squares, and triangles) as well as ten distinct colors.

### AbLang-PDB training

The AbLang-PDB model was trained using the architecture described in the Model Architecture section, utilizing the curated structural antibody dataset. During training, we maintained the pretrained weights of the base model, modifying only the QLORA adaptation parameters and the six “mixing” layers responsible for cross-chain information integration. Training employed the AdamW optimizer with a learning rate of 1e-5 and mean squared error loss function, using a batch size of 16.

To address class imbalance in the training data, we implemented a balanced sampling strategy where each epoch processed 15,270 antibody pairs, evenly distributed across three categories: overlapping epitopes, non-overlapping epitopes within the same protein family, and pairs targeting different protein families. While this approach ensured equal representation of each category during training, it resulted in more unique pairs from the non-overlapping epitope classes being trained on.

Training proceeded for 500 epochs on an NVIDIA A6000 GPU, requiring approximately 36 hours including inter-epoch evaluations. Model selection was based on ROC-AUC performance comparing training and test sets, with the epoch 240 checkpoint achieving optimal performance.

### Receiver Operating Characteristic, Precision-Recall, and F1 Score calculation

Model performance was evaluated using multiple complementary metrics implemented through scikit-learn. For receiver operating characteristic (ROC) analysis, we calculated true positive and false positive rates across 2,001 equally spaced thresholds spanning the range of possible prediction values (cosine similarity from-1 to 1 for model predictions; 0 to 1 for sequence identity comparisons). The area under the ROC curve was computed using scikit-learn’s trapezoidal rule implementation. For AbLang-RBD, ROC-AUC values were calculated separately for each of the 12 epitope classes and combined using a weighted average based on class size. For AbLang-PDB, the calculation used a binary classification scheme where overlapping epitope pairs constituted the positive class and non-overlapping pairs the negative class.

Precision-recall characteristics were assessed using scikit-learn’s precision_recall_curve and average_precision_score functions. For F1-score calculations, we utilized the previously determined optimal threshold that maximized balanced accuracy. In the case of Pfam classification, F1-scores were calculated considering all antibody pairs targeting the same protein family as positives, regardless of their specific epitope overlap status.

### LIBRA-seq dataset curation

LIBRA-seq dataset curation was performed on 7,056 class-switched antibody sequences compiled from previous LIBRA-seq experiments using PBMCs from persons living with HIV-1. Analysis included only functional, single-cell records from 10X Genomics VDJ sequencing where cells had undergone fluorescence-activated cell sorting using PE-labeled antigens, including at least one HIV envelope protein and one unrelated control antigen. Each antibody was assigned a unique identifier containing a 4-digit sequencing run prefix, with most run prefixes corresponding to unique donors except for runs 2723 and 3514, which both originated from Donor 45 (source of VRC01) ^46,64^. Nucleotide sequences were processed through IMGT HighV-Quest to determine amino acid sequences, germline gene assignments, CDR3 sequences, and percent identity to germline ^65–68^. The resulting amino acid sequences were embedded using the AbLang-PDB model, and cosine similarities were calculated between each antibody and 8ANC195. Selection of the top 20 candidates was performed blind to all functional annotations and specificity data, including suspected antigen-specificity towards positive control and negative control antigens, enabling unbiased identification of antibodies with potential epitope overlap based solely on sequence features learned by the AbLang-PDB model.

### Antibody production

Antibody heavy and light chains were synthesized as cDNA by Twist Bioscience or Genscript. Variable genes were inserted into either bicistronic plasmids encoding the constant regions of the H chain and either the kappa or lambda light chain or into separate heavy and light chain plasmids. mAbs made in house were transiently expressed using the Expifectamine transfection reagent (Thermo Fischer Scientific) in Expi293F cells in FreeStyle F17 media supplemented with 0.1% Poloxamer 188 and 20% 4mM L-glutamine (Thermo Fisher). Transfected cultures were incubated shaking for 5 days at 37°C with 8% CO2 saturation. After five days, cultures were harvested and centrifuged at a minimum of 4000 rpm for 20 minutes. Supernatant was then filtered with Nalgene Rapid Flow Disposable Filter Units with PES membrane (0.45 or 0.22 μm). Filtrate was run over PBS equilibrated columns containing protein A resin. Columns were then washed with PBS, and purified antibodies were eluted using 10mls of 100mM glycine HCL at pH 2.7 into 1 mL of 1M Tris-HCl, pH 8. These were then buffer exchanged into PBS. Remaining mAbs were synthesized by Genscript in their 10mL TurboCHO High Throughput Antibody Expression system.

### Antigen production

HPIV3 prefusion stabilized F ectodomain (Protein Data Bank [PDB] accession no. 6MJZ) was expressed in Expi293F cells through transient transfection using Expifectamine transfection reagents (Thermo Fisher Scientific) in Freestyle F17 expression media (Thermo Fisher) with the addition of 0.1% pluronic acid F-68 and 20% 4 mM l-glutamine ^69^.

Upon transfection, cultures were grown at 37 °C and 8% CO_2_ saturation levels. Six days after transfection, cultures were centrifuged at 4000 xg for 20 minutes and filtered with Nalgene Rapid-Flow Disposable Filter Units with PES membrane (0.45 or 0.22 µM).

Protein was purified through nickel affinity chromatography using an equilibrated, 1mL, prepacked HisTrap HP Column (GE Healthcare, Chicago, IL). The column was equilibriated with 15mL binding buffer (20mM sodium phosphate, 0.5M NaCl, 0.3 M imidazole, pH 7.4). Purified protein was eluted from the column with 15mL binding buffer supplemented with 0.5 M imidazole. Concentrated protein was buffer exchanged into PBS. The HisTrap purified protein was further purified by size exclusion on Superose 6 Increase 10/300 GL on the AKTA fast protein liquid chromatography (FPLC) system.

Fractions containing pure trimeric HPIV3 were identified through SDS-PAGE and the molecular mass. Antigenicity was confirmed with binding to 3X1. Protein concentration was quantified using UV/visible spectroscopy and frozen at-80°C until use.

HIV-1 envelope proteins (BG505, CZA97, and ZM106.9) were designed using the SOSIP platform to yield soluble Env proteins stabilized in the pre-fusion conformation. These SOSIP constructs incorporated several stabilizing mutations: an intermolecular disulfide bond between gp120 and gp41 (A501C and T605C), a trimer-stabilizing mutation (I559P), a truncated gp41 transmembrane region at position 664, and an I201C/A433C mutation to inhibit CD4-induced movement of Env. Additionally, a flexible serine-glycine linker was inserted between gp120 and gp41 (positions 507 and 512) to create single-chain constructs ^70^.

HIV-1 envelope proteins were expressed in a highly similar fashion but with the following caveats. Post-culture and centrifugation, the filtered supernatant was applied to an affinity column of agarose-bound Galanthus nivalis lectin (Vector Laboratories) at 4°C. After washing with PBS, proteins were eluted with 30 mL of 1 M methyl-α-D-mannopyranoside. The eluate was buffer-exchanged three times into PBS and concentrated using either 30 kDa or 100 kDa Amicon Ultra centrifugal filter units.

Final purification was achieved by size-exclusion chromatography using either a Superose 6 Increase 10/300 GL or Superdex 200 Increase 10/300 GL column on an AKTA FPLC system. Fractions corresponding to correctly folded trimeric Env proteins were collected and validated by SDS-PAGE for molecular weight determination and by ELISA for antigenicity using Env-specific monoclonal antibodies.

### Indirect ELISA

In a 96-well plate, 100 μL of antigen was coated at 2 ug/mL overnight at 4°C. The plates were then washed three times with PBS supplemented with 0.05% Tween20 (PBS-T) and blocked using 5% bovine serum albumin in PBS. Plates were incubated for one hour at room temperature and then washed three times using PBS-T. Primary antibodies were diluted in 1% BSA in PBS-T starting at 10 μg/mL with a 1:5 dilution.

After incubating at room temperature for one hour and washing with PBS-T, 100 μL of goat anti-human IgG conjugated to peroxidase was added at a 1:10,000 dilution in 1% BSA in PBS-T. These were incubated for one hour at room temperature, washed three times with PBS-T, then developed using TMB substrate. Plates developed for ten minutes at room temperature and were then stopped using 1 N sulfuric acid. Then, absorbance was then measured at 450 nm.

### Competition ELISA

Wells of a 96-well plate were coated with 100 μL of 2 μg/mL purified BG505 N332T SOSIP and were left at 4 °C overnight. Plates were then washed three times using PBS-T and each well was blocked using 100ul of 5% BSA in PBS for 1 hour. After washing three times using PBS-T, primary antibodies were diluted 10-fold starting at 100 μg/mL using 1% BSA in PBS-T and 75 μL was added each well. After incubating for one hour at room temperature, without washing, 25 μL of biotinylated antibody was added to each well to the final concentrations of 1 μg/mL and 0.1 μg/mL. This was incubated at room temperature for one hour and was washed three times using PBS-T. Then, 100 μL of streptavidin-HRP at a dilution of 1:10,000 in 1% BSA in PBST was added to each well and was incubated for one hour at room temperature. These plates were then washed three times and bound antibodies were detected using TMB substrate and sulfuric acid. Competition ELISAs were repeated at least 2 times. Data is displayed as the percentage change in binding relative to the binding of an antibody when no competitor is present.

### Structural representation of HIV reference antibodies G

A composite image of VRC01, 8ANC195, and PG9 binding BG505 Envelope was generated by first loading PDB ID 5VJ6 into open-source PyMOL™ © Schrodinger, LLC Version 2.4.0 ^49,55^. The antibody-antigen complex was represented as a surface, and PG9 was colored wheat, 8ANC195 in light green, gp120 in light gray, and gp41 in dark gray. PDB ID 8VGW was then loaded into PyMOL and the gp120 structures from one protomer were aligned to that of one gp120 protomer in 5VJ6. VRC01 was then colored pink and shown as a surface without visualization of the Envelope protein present in its native complex. Finally, ray tracing was performed with default parameters.

## Conflicts of Interest

I.S.G. is listed as an inventor on patents filed describing antibodies characterized here.

I.S.G. is listed as an inventor on the patent applications for the LIBRA-seq technology.

I.S.G. is a co-founder of AbSeek Bio. I.S.G. has served as a consultant for Sanofi. The Georgiev laboratory at VUMC has received unrelated funding from Merck and Takeda Pharmaceuticals.

## Notes

https://github.com/IGlab-VUMC/AbLangRBD1

https://github.com/IGlab-VUMC/AbLangPDB1

### Summary of Updates

Acknowledgements section updated to better reflect funding.

