## Supplementary figures and images for "Contrastive Learning Enables Epitope Overlap Predictions for Targeted Antibody Discovery"

### Supplemental Figures

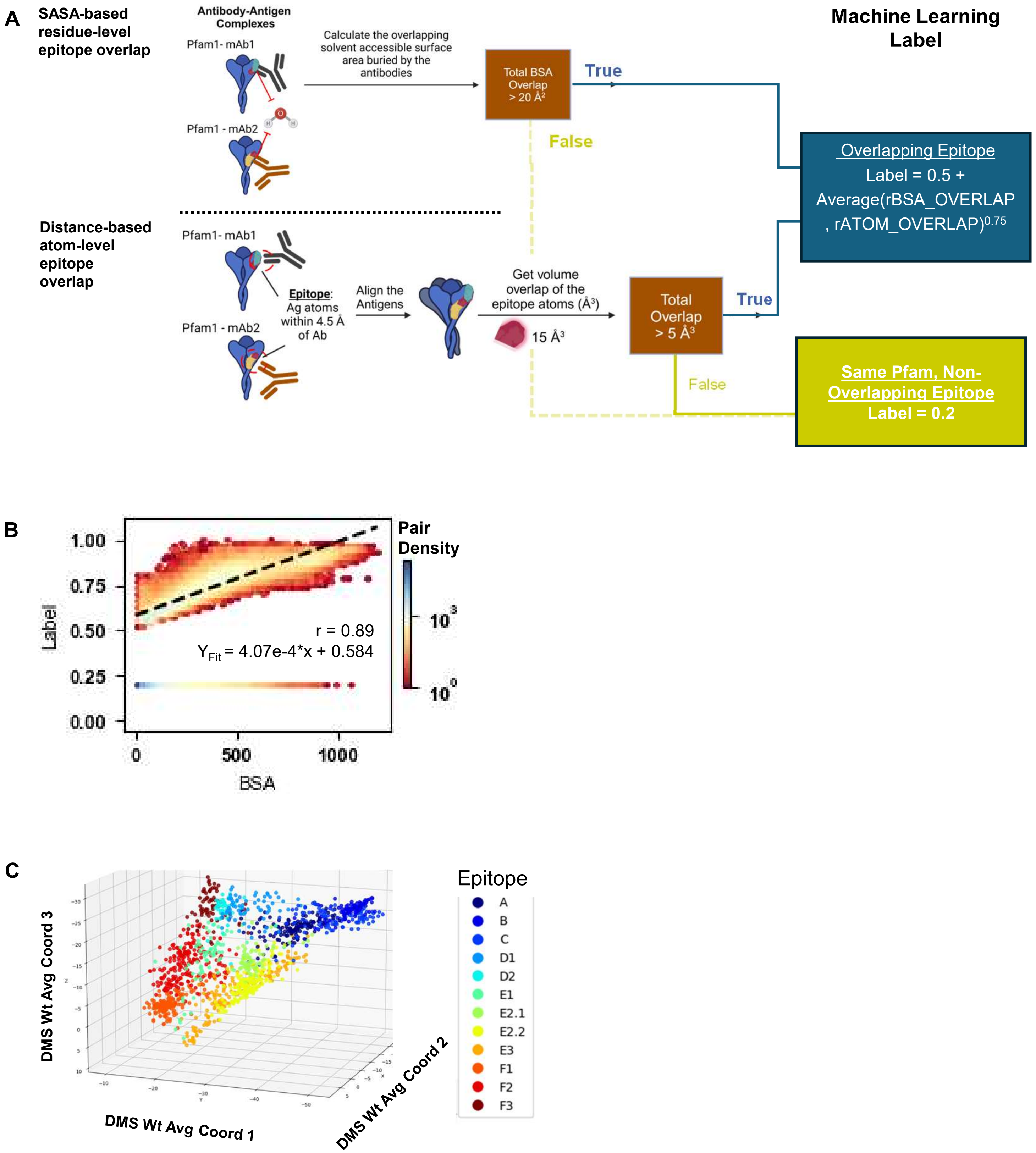

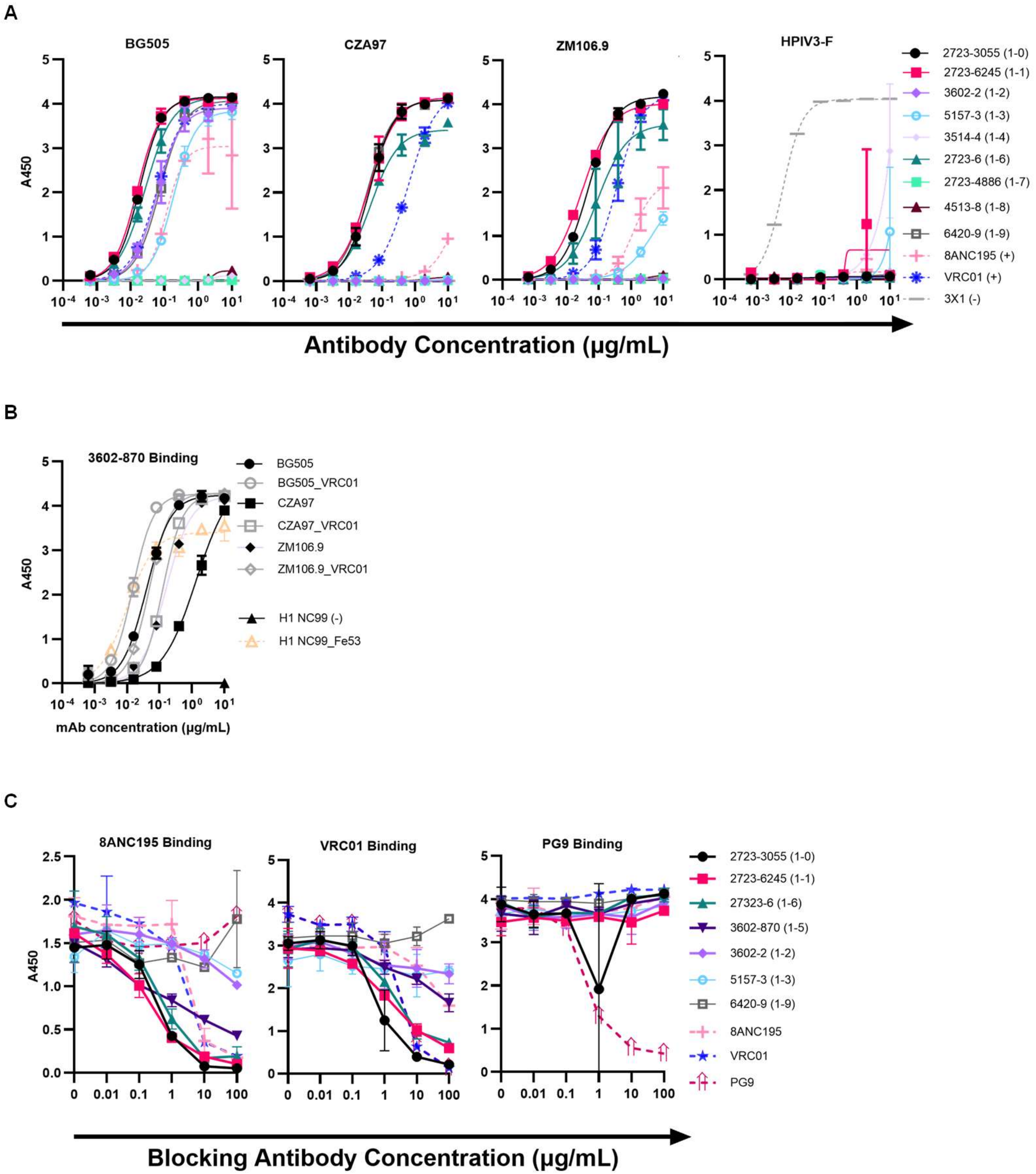
